# Adaptation optimizes sensory encoding of future stimuli

**DOI:** 10.1101/2024.03.20.585768

**Authors:** Jiang Mao, Constantin Rothkopf, Alan A. Stocker

**Author notes:** Correspondence: Dr. Alan A. Stocker, Computational Perception and Cognition Laboratory, Goddard laboratories, Rm 421, 3710 Hamilton walk, Philadelphia, PA 19106, U.S.A.

## Abstract

Sensory neurons continually adapt their response characteristics according to recent sensory input. However, it is unclear how such a reactive process shaped by sensory history can benefit the organism going forward. Here, we test the hypothesis that adaptation indeed acts proactively in the sense that it optimally adjusts sensory encoding for the future, i.e. for the next expected sensory input. We first quantified adaptation induced changes in sensory encoding by psychophysically measuring discrimination thresholds for visual orientation under different adaptation conditions. Using an information theoretic analysis, we found that adaptation consistently reallocates coding resources such that encoding accuracy peaks at the adaptor orientation while total coding capacity remains constant. We then asked whether this characteristic change in encoding accuracy is predicted by the temporal statistics of natural visual input. By analyzing the retinal input of freely behaving human subjects in natural environments, we found that the distribution of local visual orientations in the retinal input stream at any moment in time is also peaked at the mean orientation computed over a short input history leading up to that moment. We further tested our hypothesis with a recurrent neural network trained to predict the next frame of natural scene videos (PredNet). We simulated our human adaptation experiment with PredNet while analyzing its internal sensory representation. We found that the network exhibited the same change in encoding accuracy as observed in human subjects, and as predicted by the natural input statistics. Taken together, our results suggest that adaptation induced changes in encoding accuracy are an attempt of the visual systems to be best possibly prepared for future sensory input.

## Introduction

Biological information processing systems continually adapt their sensory representations to statistical changes in their sensory environment. This is well documented by the various changes in neural response characteristics that occur after prolonged exposure to an adaptor stimulus such as a fixed visual orientation (e.g., Dragoi et al., 2000, 2002; Patterson et al., 2013) or motion direction (e.g. Kohn and Movshon, 2003, 2004), as well as the corresponding changes at the perceptual level (e.g. Regan and Beverley, 1985; Clifford et al., 2001; Mitchell and Muir, 1976). Many popular visual illusions represent particularly salient examples of how adaptation can affect perception (e.g., the motion aftereffect (Mather et al., 1998)). Adaptation manifests itself at every level of information processing, affecting every neuron involved in the representation and processing of sensory information, and the effects accumulate and interact along the hierarchically organized sensory representations in the brain (e.g., Kohn and Movshon (2004); Stocker and Simoncelli (2009)). Its ubiquitous nature suggests that adaptation provides fundamental and important benefits that must clearly outweigh the costs of creating perceptual distortions.

Adaptation has been thought of as a mechanism that adjusts neural representations to the spatiotemporal statistics of the current sensory context in order to maximize the amount of information encoded about the sensory input (Barlow et al., 1961; Wainwright, 1999; Brenner et al., 2000b; Sharpee et al., 2014). This efficient coding hypothesis has been empirically validated at the level of single neurons in simple systems where input-output relations can be relatively easily controlled and manipulated (e.g. the motion sensitive neurons of the blowfly (Brenner et al., 2000a; Fairhall et al., 2001)). In more complex neural systems such as the primate brain, perceptual variables are typically encoded in a distributed fashion over an entire neural population. In this case, the efficient coding hypothesis is more difficult to test because it requires sufficiently comprehensive measurements of the neural code to faithfully calculate the mutual information content across the entire neural population.

Approximating mutual information with Fisher information (Fisher, 1922) can help resolving these difficulties as Fisher information can be more directly computed and related to neurophysiological parameters (Seung and Sompolinsky, 1993; Brunel and Nadal, 1998). Fisher information is a local measure of sensory encoding accuracy, and can be interpreted as the amount of coding resources allocated to represent a particular stimulus value (Wei and Stocker, 2015). Furthermore, Fisher information provides a lower bound on perceptual discriminability (Seung and Sompolinsky, 1993; Seriès et al., 2009; Zhang and Stocker, 2022). As a result, adaptation induced changes in sensory encoding can be directly quantified with psychophysical measurements of discrimination thresholds (in units of Fisher information).

Previous studies have measured the effect of adaptation on discriminability but rarely did so for the entire stimulus range, which is necessary to quantify changes in the distribution of Fisher information. Some studies have shown that adaptation decreases discrimination threshold at the adaptor but increases the threshold for stimulus values near but different from the adaptor (Regan and Beverley, 1983, 1985; Clifford et al., 2001; Phinney et al., 1997). For circular variables like orientation, some studies found that discrimination thresholds also decrease for stimulus values that are opposite of the adaptor (i.e. for orientation, orthogonal to the adaptor) (Clifford et al., 2001; Dragoi et al., 2002), but this result is not conclusive (Westheimer and Gee, 2002; Clifford et al., 2003). Thus, currently available discrimination data are not sufficient to fully characterize adaptation induced changes in sensory encoding. What is missing are thresholds measurement across the entire stimulus range for a given, identical adaptation state.

Testing the efficient coding hypothesis of adaptation also requires knowledge of the contextual stimulus distribution for which the representation is optimized for. For efficient codes that maximize mutual information, Fisher information should be proportional to the square of this distribution (Wei and Stocker, 2015, 2016, 2017). Previous studies have shown that under quasi-stationary contexts (i.e., long timescales) sensory encoding is qualitatively well matched to the overall statistics of the observer’s natural environment. For example, the distribution of local orientations in natural visual scenes shows strong peaks at cardinal orientations (Coppola et al., 1998) that are well aligned with the reported higher orientation discriminability of human observers at cardinal orientations (Caelli et al., 1983; Ganguli and Simoncelli, 2010). However, for the dynamic contexts that are relevant for adaptation (i.e., short timescales), these input distributions are more difficult to define and measure. In vision particularly, the observer’s active control of the gaze position can strongly affect the shape of the input distributions at the level of the retina (Rothkopf et al., 2009; Rothkopf and Ballard, 2009; Straub and Rothkopf, 2021). Thus input distributions over the timescales relevant for adaptation must be determined based on the retinal image under natural viewing conditions.

Conceptually, the efficient coding hypothesis of adaptation seems somewhat at odds with the notion that adaptation is driven by the recent stimulus history; to be truly useful for the observer, sensory representations should be optimized for future rather than past sensory input. Of course, it is well established that due to its continuous nature, the recent state of our environment is a good predictor of its future (Dragoi et al., 2002; Felsen et al., 2005; van Bergen and Jehee, 2019). However, again this does not necessarily translate to the input distributions at the level of the retina due to the observer’s active control and selection of sensory input via eye-movements (Yarbus, 1967; Findlay and Gilchrist, 2003; Hayhoe and Ballard, 2005; Land and Tatler, 2009; Najemnik and Geisler, 2005).

In this study, we investigate the efficient coding hypothesis of adaptation for visual orientation perception in human observers. We first psychophysically characterize changes in orientation discrimination thresholds across the entire orientation range induced by prolonged exposures to different adaptor stimuli with different orientations. We show that while adaptation decreases discrimination thresholds at the adaptor orientation and - surprisingly - also at orientations orthogonal to the adaptor, it increases discrimination thresholds further away from the adaptor orientation. Adaptation thereby does not change total Fisher information (i.e., the overall amount of coding resource) but rather reallocates coding resources, which can be expressed with a single, isomorphic adaptation kernel for all adaptor orientations. We then show that this kernel matches the spatiotemporal orientation distributions in retinal input of human observers freely behaving in natural outdoor environments. After relatively stable visual input during a short time-window, the distribution of local orientations in the retinal image at the next time step is sharply peaked at the mean orientation over the time-window. Finally, we use an artificial recurrent neural network, designed and trained to predict the next frame of naturalistic videos (Lotter et al., 2016), to demonstrate that the adaptation kernel we extracted from human psychophysics and natural scene statistics is indeed beneficial for the perception of the very next visual input. When presented with the same adaptation stimulus sequence used in our human psychophysical experiment, the network’s internal sensory representations developed a very similar reallocation of coding resources as observed in human subjects.

Taken together, our results suggest that adaptation maintains an efficient encoding of stimulus features given the spatiotemporal statistic of sensory signals in natural environments, aimed at providing a best possible representation of the next sensory input.

## Results

The encoding capacity of a sensory system is limited, and is usually not uniformly distributed over the full range of a stimulus variable. We hypothesize that adaptation temporarily reallocates this encoding capacity without changing the total amount of representational resource. In the following, we first formalize this reallocation process with a model before we test it empirically.

### Reallocation Model

Consider the sensory representation of local orientation in the visual system. We use Fisher information *J* (*θ*) to quantify the coding resource allocated to encode visual orientation. For quasistationary conditions (i.e., over long timescales) in natural visual environments, we assume that Fisher information is higher around cardinal orientations (Fig. 1). This assumption is qualitatively aligned with the observed increase in discriminability at cardinal compared to oblique orientations (Caelli et al., 1983), as well as with the idea of efficient coding given the overall statistics of local visual orientations in natural scenes (Coppola et al., 1998). Applying a transformation 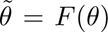 where *F* (*θ*) is the cumulative distribution of the square-root of the Fisher information maps the representation in stimulus space to a sensory space where the distribution of Fisher information is uniform (Wei and Stocker, 2012, 2015). *F* (*θ*) can be thought of as the transformation by which the stimulus information is mapped onto an internal (i.e., neural) sensory representation. If encoding is efficient and aimed at maximizing mutual information between stimulus and encoded value, then *F* (*θ*) is identical to the cumulative function of the overall stimulus distribution *p*(*θ*) (i.e., the prior), giving rise to the efficient coding constraint (Wei and Stocker, 2015, 2016, 2017)

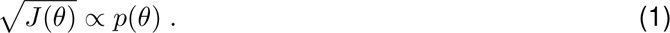

**Figure 1:**
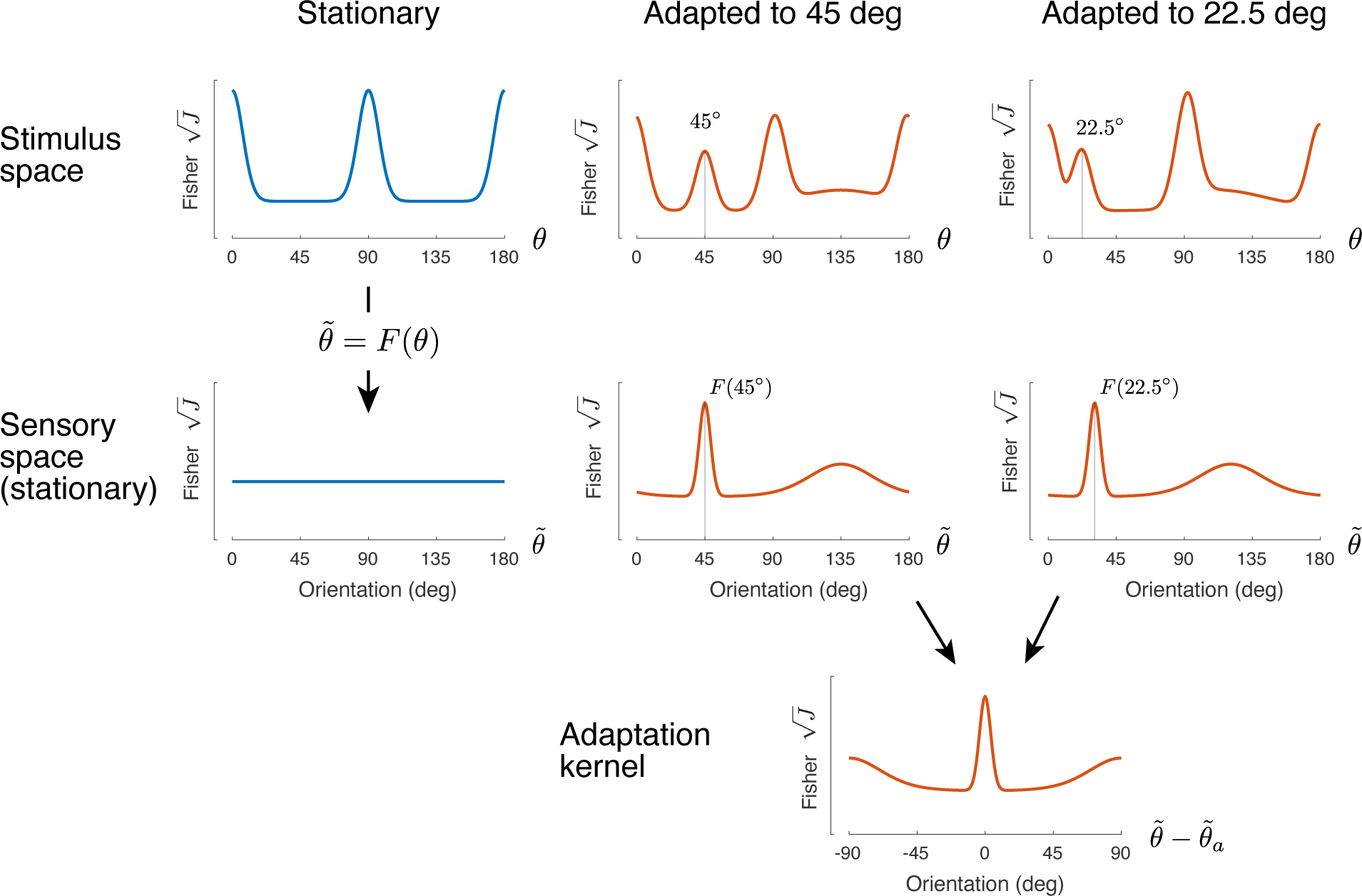
Reallocation Model. Under (quasi-)stationary conditions (left), Fisher information *J* (*θ*) has a certain distribution across the entire stimulus range (e.g., orientation), which transforms to a uniform distribution in the sensory space *F* (*θ*). After adaptation (middle: adaptor at 45 deg, right: adaptor at 22.5 deg), Fisher information is reallocated according to an isomorphic adaptation kernel, which is identical in shape for different adaptors but centered at the respective adaptor orientation. Total Fisher information remains constant across all conditions.

The reallocation model assumes that after adaptation to a stimulus containing a single orientation, Fisher information is temporarily redistributed according to an isomorphic adaptation kernel in sensory space, i.e., the reallocation of sensory coding resources follows a fixed pattern relative to the orientation of the adaptor stimulus that is independent of the adator orientation. The adaptation kernel is expressed in units of square-root of Fisher information (Fig. 1). If encoding is efficient (Eq. (1)), the kernel should reflect the changes in stimulus distribution (i.e., the prior) for which adaptation is meant to optimize the code.

After adaptation, the distribution of Fisher information in sensory space is no longer uniform. However, we can again transform sensory space to a post-adaptation state where the distribution of Fisher information is once more uniform using a second, adaptation induced transformation that now represents the cumulative of the adaptation kernel. With the assumption that the noise in sensory space is homogeneous when Fisher information is uniform, we can directly extract Fisher information and its changes under different adaptation conditions (i.e., the adaptation kernel) from psychophysical measurements of discrimination threshold. Likewise, we can predict the psychometric functions of the discrimination experiment that correspond to a given adaptation kernel (see Methods for details).

### Psychophysical experiment

Five human subjects performed a two alternative forced-choice (2AFC) orientation discrimination experiment under different adaptation conditions (Fig. 2). At the beginning of each block, subjects were presented with an adaptor stimulus for one minute. After that, subjects performed 192 trials of the 2AFC task, reporting which one of two orientated grating stimuli was more clockwise/counter-clockwise. Every trial started with a 5s period of top-up adaptation. Each adaptor consisted of 8 blocks.

**Figure 2:**
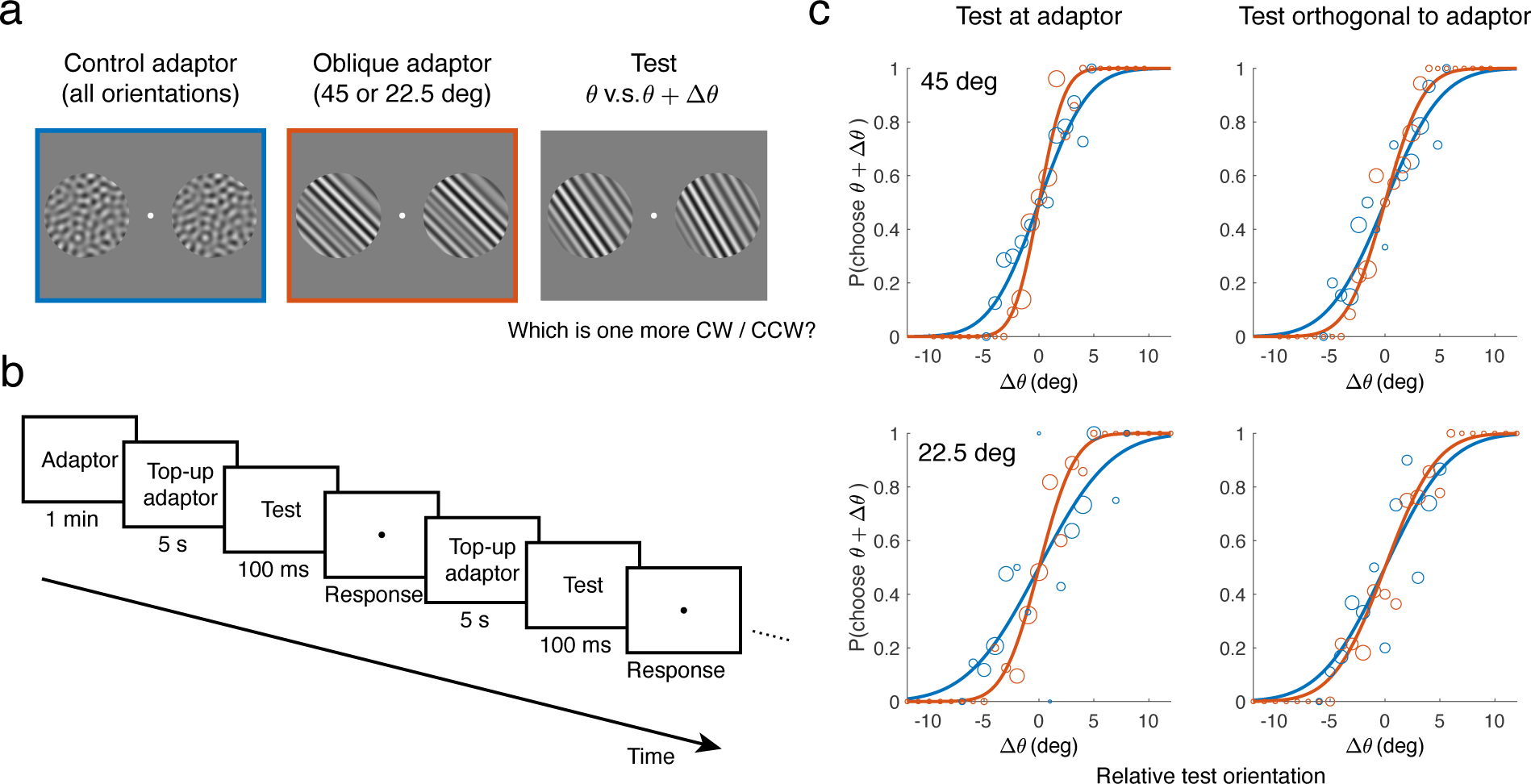
Experimental procedure and measured adaptation induced changes in discrimination thresholds. (a) Stimuli were bandpass filtered white noise with either a uniform (control adaptor) or narrowband orientation spectrum (oblique adaptor, test stimuli). (b) Task structure (single block). At the beginning of each block, there was an adaptation period of 1 minute. Every trial started with 5s top-up adaptation, after which subjects were briefly presented with two test stimuli and asked to report which one was more clockwise/counter-clockwise. Subjects performed the same number of trials for each of the three adaptor conditions (two oblique and the control adaptors). (c) Data and fitted psychometric curves at two test orientations of one example subject (Subject 1). Adaption to the oblique adaptors (orange) results in steeper psychometric curves for test orientations at and orthogonal to the adaptor orientation compared to the control adaptor condition (blue). The size of data points is proportional to the number of trials at that test orientation.

We tested two types of adaptors. The oblique adaptor is identical in structure to the test stimuli (i.e., same bandpass and orientation filter), with the orientation filter centered either at 45 deg or 22.5 deg (0 deg being vertical). We chose oblique as opposed to cardinal adaptor orientations to avoid possible ceiling effects; discrimination thresholds are known to be lowest at cardinal orientations (Caelli et al., 1983) and adaptation is expected to lower it further at the adaptor orientation (Regan and Beverley, 1985; Clifford et al., 2001). The second, control adaptor is identical to the oblique adaptor excepted that it has a uniform orientation filter profile. Importantly, discrimination thresholds measured under this well-defined, control adaptation state serve as a reference against which we compare changes in discrimination threshold induced by the oblique adaptor (see Methods).

We fit psychometric curves to the 2AFC data (Fig. 2c), and extracted discrimination thresholds (Fig. 3). In the control adaptation condition, discrimination thresholds were lowest at cardinal orientations as shown in previous studies without adaptation (Caelli et al., 1983). Adaptation to an oblique adaptor causes discrimination thresholds to decrease at but increase slightly away from the adaptor orientation compared to the control condition. This is consistent with previous findings (Regan and Beverley, 1985; Clifford et al., 2001). Interestingly, discrimination thresholds at test orientations orthogonal to the adaptor also decreased after adaptation, which has not been consistently shown (Clifford et al., 2001; Westheimer and Gee, 2002; Clifford et al., 2003; Dragoi et al., 2002). These general results hold for both oblique adaptors (45 deg and 22.5 deg) and across all subjects (Fig. 4).

**Figure 3:**
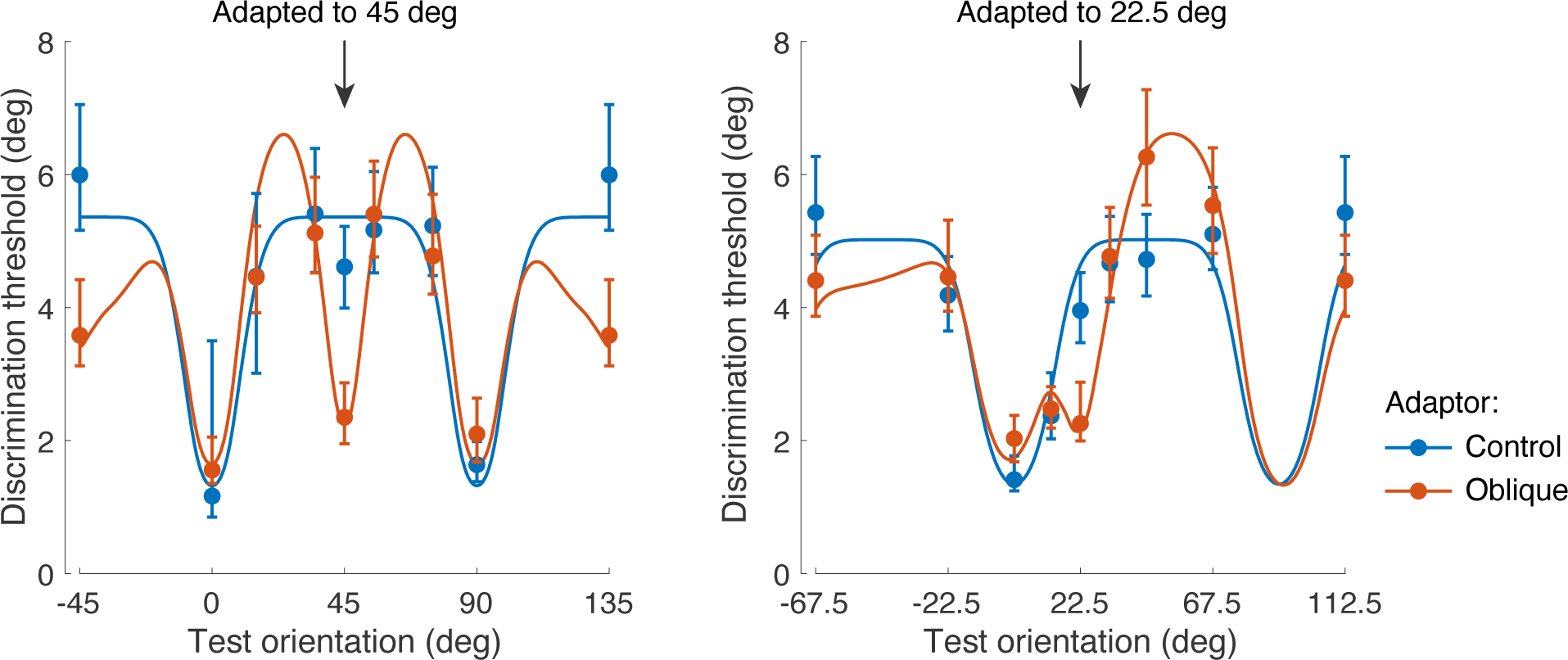
Discrimination threshold data and model fits averaged across subjects (see Fig. 4). With the control adaptor, thresholds are lower at cardinal orientations. With the oblique adaptors, thresholds decrease at and orthogonal to the adaptor, and increase away from the adaptor. Error bars represent the 95% intervals computed over 1000 bootstrap samples of the data (individual subjects).

**Figure 4:**
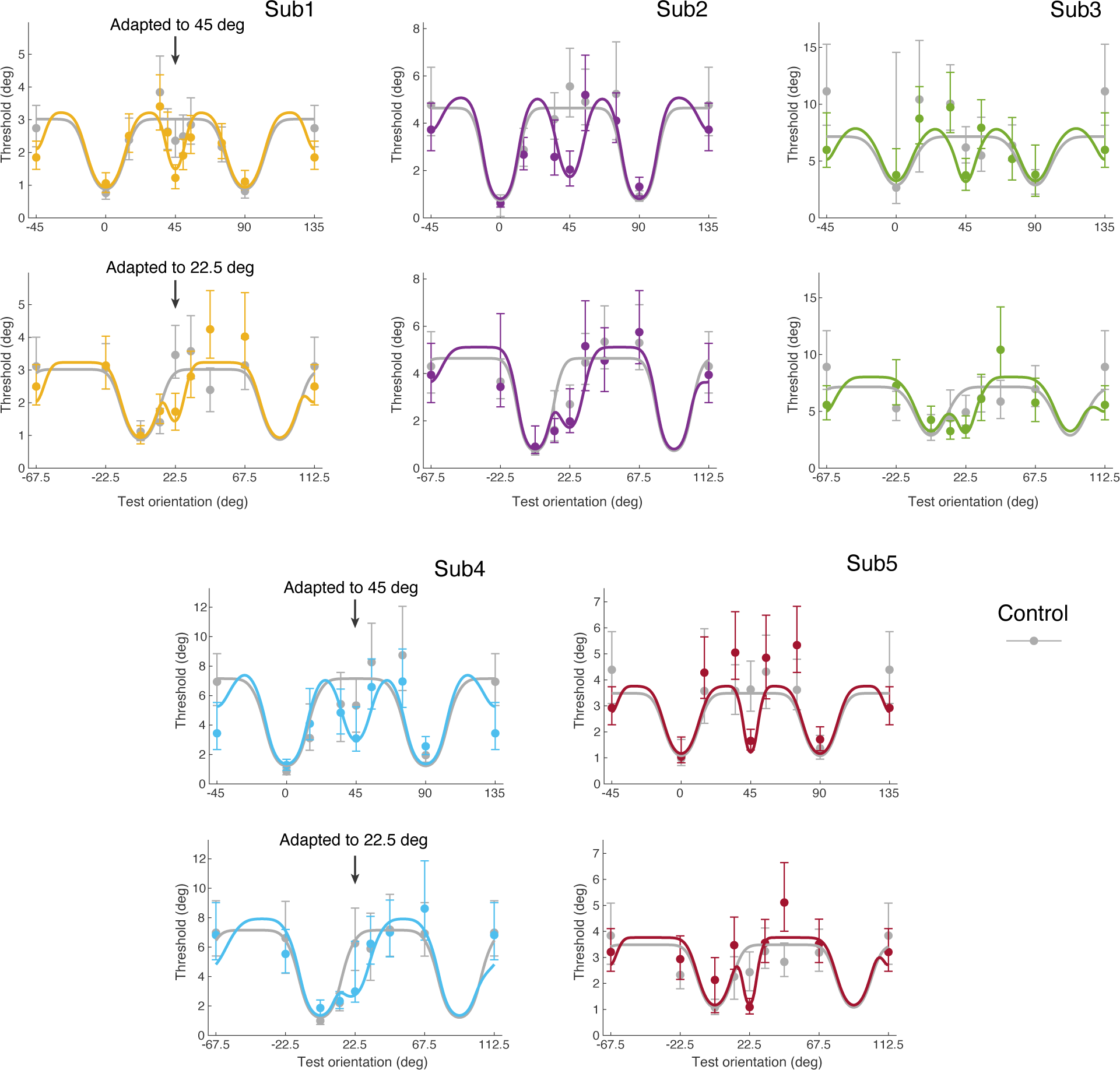
Discrimination threshold data and model fits for individual subjects. Compared to the control condition, subjects consistently show lower thresholds at and orthogonal to the adaptor after adaptation, and higher thresholds slightly away from adaptor. Error bars represent the 95% intervals over 1000 bootstrap samples of the data.

### Model Fit and Comparison

As our experimental results show, discriminability improves after adaptation for test orientations both at and orthogonal to the adaptor orientation. We thus assume an adaptation kernel that peaks at and orthogonal to the adaptor orientation (Fig. 1). We individually fit the data from each subject with the reallocation model. First, we fit the data measured under the control condition, which allowed us to determine the distribution of Fisher information before adaptation (i.e., the subject’s individual stationary sensory space 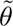) and the overall amount of coding resource available (i.e., the subject’s total Fisher information, which is inversely proportional to the noise variance in the sensory space). We then jointly fit the data from the 45 deg and 22.5 deg adaptor conditions and determined the adaptation kernel of every subject. Figure 3 shows the average measured and predicted discrimination threshold based on model fits to individual subjects’ data. Figure 4 shows the measured and predicted thresholds of individual subjects. The model captures not only the improvement in discriminability at and orthogonal to the adaptor orientation, but also the increases in discrimination threshold at test orientations slightly different from the adaptor orientations.

Figure 5a shows the extracted adaptation kernels for every subject. The kernels are similar across subjects, consistently showing a sharp peak in Fisher information at the adaptor as well as a more shallow peak for orientations orthogonal to the adaptor. Note that the kernels are plotted in each subject’s individual sensory space 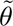.

**Figure 5:**
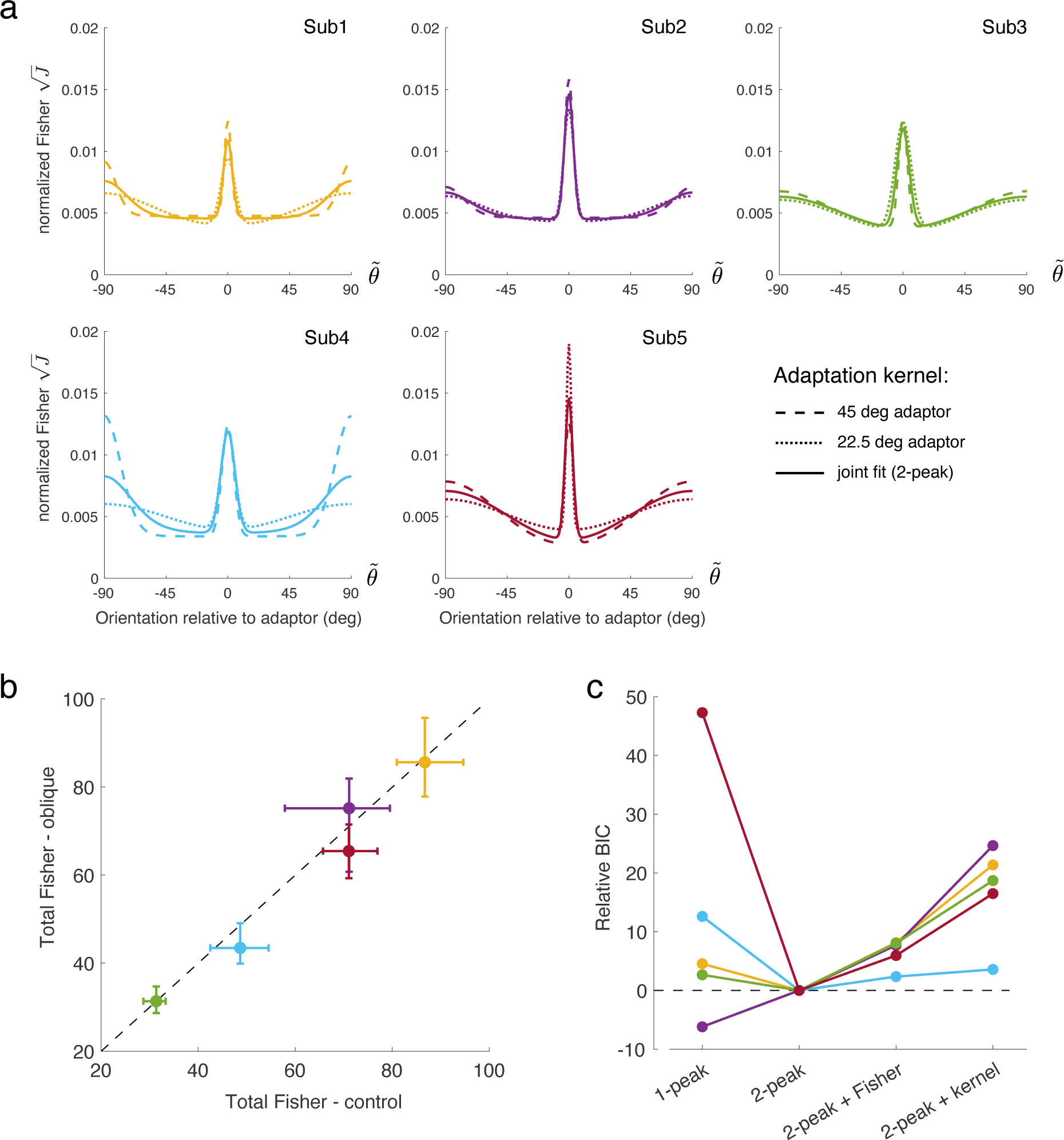
Model comparison. (a) Extracted adaptation kernels. Kernels are similar across subjects and do not substantially differ when independently fit to data from the two oblique adaptor conditions. (b) Total Fisher information (i.e. total coding resource) extracted from separate fits to control and oblique adaptor conditions. Values vary across subjects but fall close to the unity line. (c) Testing the key assumptions of the reallocation model using a BIC goodness-of-fit criterion: original model (2-peak), relaxing the fixed resource assumption (2-peak + Fisher), relaxing the single kernel assumption (2-peak + kernel), and assuming a kernel with only a peak at the adaptor (1-peak). Error-bars in all panels represent 95% confidence intervals from 100 bootstrap samples of the data. Subject color code as in Fig. 4.

We can further test key assumptions of the proposed reallocation model. For example, a separate fit of the data obtained in the two oblique adaptor conditions results in adaptation kernels that do not differ much from the kernels obtained from a joint fit, suggesting that adaptation is indeed governed by a mechanism that is independent of the specific adaptor orientation (Fig. 5a). Also, when fit separately, the fit total Fisher information for each condition is very similar, thus confirming our assumption that the total coding resource remains fixed and does not change with the adaptation state (Fig. 5b). Finally, we performed a formal model test where we compared the proposed real-location model (2-peak) with a model that assumes coding improvement only at but not orthogonal to the adaptor (1-peak), a model that allows the total Fisher information to change with adaptation (2-peak + Fisher), and a model that allows the adaptation kernel to be different for different adaptor orientations (2-peak + kernel). When appropriately penalized for the number of free parameters (BIC), the reallocation model (2-peak) best fits the data for all subjects except Subject 2 for whom the only moderate threshold improvement orthogonal to the adaptor favors an adaptation kernel with a single peak at the adaptor (Fig. 5c).

### Natural Scene Statistics

Next, we analyzed the temporal statistics of visual orientations in retinal images of freely behaving human observers. In particular, we asked whether the measured adaptation kernels match the distributions of visual orientation in the retina image stream for similar temporal contexts as in our psychophysical adaptation experiments.

Subjects took a stroll through a forest while wearing an SMI ETG2 head-mounted camera and an eye-tracker recording the scene and their eye movement, respectively (Fig. 6a). In every frame of the video, we considered a central image patch (6×6 deg visual angle around the gaze center; Fig. 6b) and extracted the local orientation at each position within the image patch based on a linear multi-scale, multi-orientation image decomposition (steerable pyramid; Simoncelli and Freeman (1995)). We computed mean and variance of orientation over a sliding 3s time-window, and then identified times and positions where the variance over the time-window was relatively small (circular variance smaller than 0.1). For those spatiotemporal positions, we calculated the difference between the orientation in the next frame and the mean orientation over the preceding 3s time-window. We found that the orientation in the next frame is mostly likely to be similar to the mean orientation in the immediate past at the same position (Fig. 6c, left). The less stable the histories are (larger variances), the wider the distributions of orientations in the next frame, until they eventually become uniform (Fig. 6c, right). This is consistent across spatial frequency levels (see Supplementary Fig. 3). These statistical regularities imply that reallocating Fisher information towards the adaptor orientation is consistent with the efficient coding hypothesis under natural viewing conditions with stable input history. Likewise, for highly variable input histories the distributions are almost uniform and therefore efficient coding does not propose any reallocation of resources; this corresponds to the control adaptor condition in our psychophysical experiment and reallocation model.

**Figure 6:**
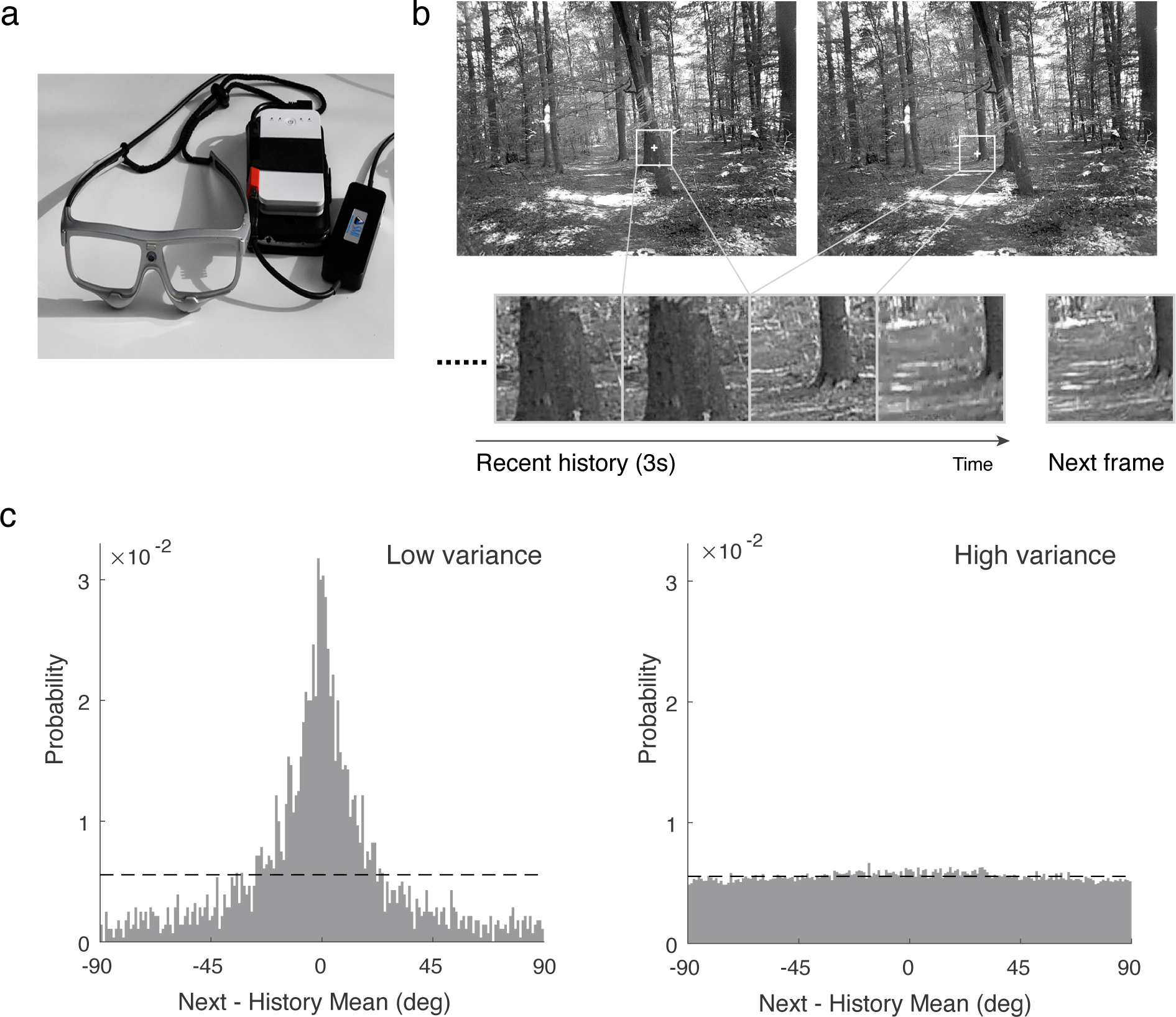
Retinal input statistics under natural viewing condition. (a) The combination of head-mounted camera and mobile eye-tracker allowed us to extract the retinal input statistics of human subjects freely behaving (i.e., walking) in a natural, forest environment. (b) Within a small window centered at the subjects’ gaze location (white frame), we computed the distribution of local visual orientation in the next frame relative to the mean orientation over an immediately preceding, three seconds long time-window (history mean; same spatial position). (c) Distribution of orientations in the next frame relative to the history mean when the variance of orientations in the time-window was low (<0.1; left) or high (>0.9; right), respectively.

### Predictive Neural Network

Previous studies have shown that deep neural networks implicitly learn to encode stimulus features as predicted by efficient coding (Benjamin et al., 2022). Here we use “PredNet”, a recurrent neural network designed and trained to predict the next frame in a video sequence (Lotter et al., 2016), to test to what degree sensory representations can dynamically change depending on the temporal input statistics. Previous studies have shown that PredNet reports illusory motions similar to those perceived by human subjects (Watanabe et al., 2018; Kobayashi et al., 2022). PredNet was inspired by the concept of “predictive coding” in neural networks (Rao and Ballard, 1999). PredNet uses top-down connections conveying the local predictions of incoming stimuli and bottom-up signals of the deviations from the predictions (Fig. 7a). There are four sub-layers in each layer *i* of PredNet: a recurrent representation layer (*R_i_*), a prediction layer (*Â_i_*), an input or target layer (*A_i_*), and an error layer (*E_i_*). The representation layer takes feedback from the error layer and the higher representation layer and generates predictions for the input layer; the error layer calculates the deviation of the prediction layer from the input layer and passes them on to the next input layer. Importantly, PredNet only contains static (i.e., non-adaptive) neurons.

**Figure 7:**
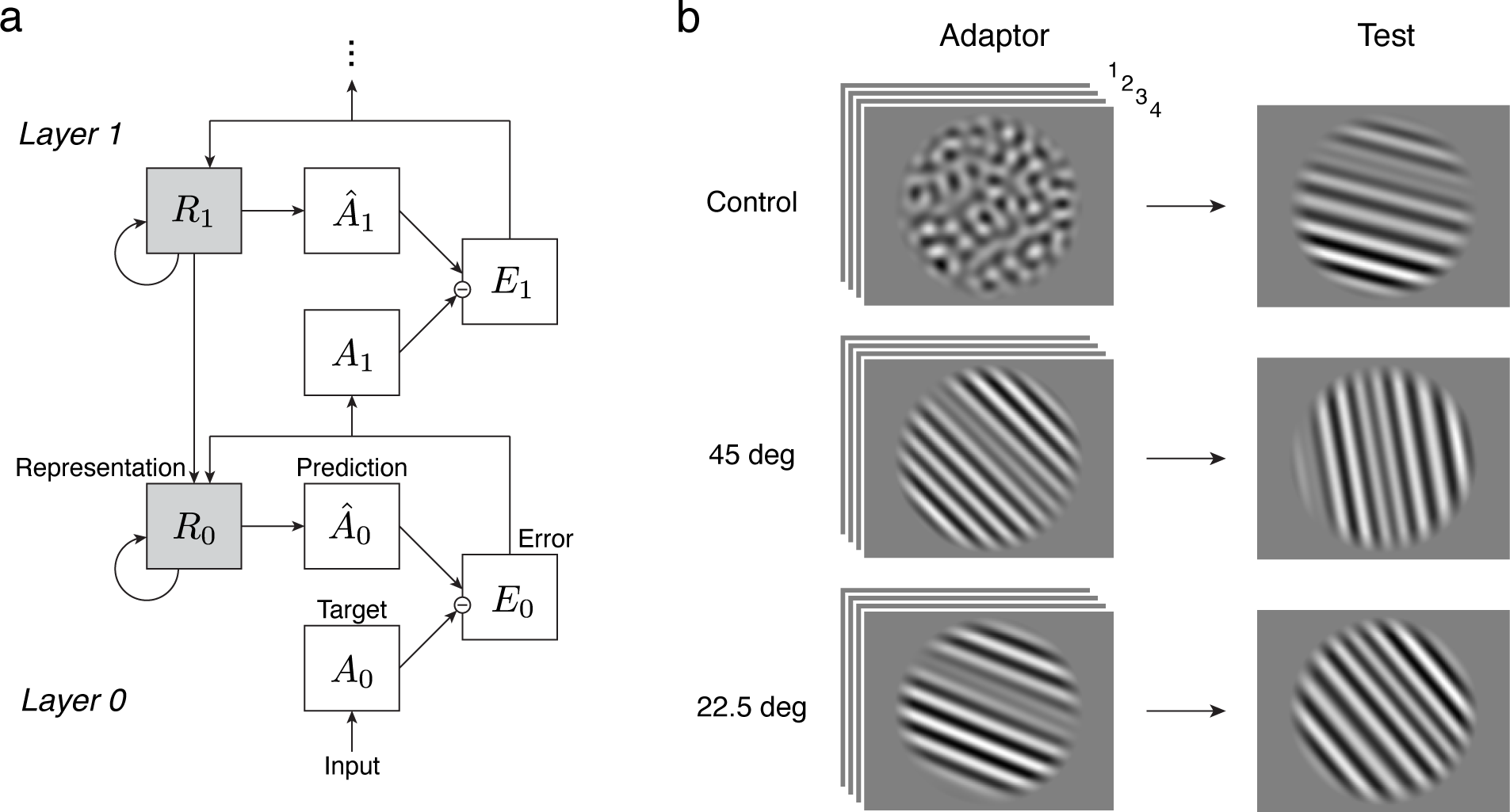
PredNet architecture and adaptation experiment in PredNet. (a) Architecture of PredNet (Lotter et al., 2016). Each layer of PredNet consists of four sub-layers: a representation layer (*R_i_*), a prediction layer (*Â_i_*), an input or target layer (*A_i_*), and an error layer (*E_i_*). The representation layer takes feedback from the error layer and the higher representation layer and makes predictions about the next input. The error layer takes the difference between the prediction and the input and passes it on to the next layer. The network has four layers in total. (b) Adaptation experiment in PredNet. In each trial, PredNet is presented with four frames of the adaptor stimulus (control and oblique adaptor), followed by one frame of the test stimulus. The stimuli were the same as in the human psychophysical experiment (Fig. 2). Encoding accuracy was measured based on the network’s activity in the lowest representational layer in response to the test stimulus.

We tested how the prolonged exposure to an adaptor stimulus affects PredNet’s encoding accuracy in its representational layers. We used PredNet pretrained on video sequences from the KITTI natural scenes data set (Geiger et al., 2013). We exposed the network to the same stimulus sequence as used in our human adaptation experiment (Fig. 7b). We input a sequence of four adaptor frames (mimicking the adaptation phase) followed by a test frame, and then computed Fisher information in the lowest representation layer in response to the test stimulus. With the assumption that independent Gaussian noise corrupts the response in each neuron, Fisher information is equivalent to the squared gradient of the neural response in the direction of the test orientation (see Methods). In this way, we computed Fisher information in the first representation layer as a function of the test orientation *θ* for all three adaptor conditions (control; 45 deg and 22.5 deg oblique adaptor).

Figure 8a shows the resulting Fisher information distributions when plotted in stimulus space. In the control adaptor condition, encoding accuracy in the first representational layer of PredNet is higher at cardinal compared to oblique orientations. After adaptation to the oblique adaptors, however, Fisher information peaks at the adaptor orientation. Both effects qualitatively match the measured changes in encoding accuracy in human subjects (Fig. 4). Furthermore, when plotted in sensory space 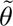 as defined by the networks’ encoding precision measured in the control adaptor condition, Fisher information well resembles the adaptation kernels extracted from human subjects (Fig. 5a). For both oblique adaptor conditions, the curves show a similar symmetric peak in encoding accuracy at the adaptor orientation and also the slight reduction in accuracy for orientations in the vicinity of the adaptor (Fig. 8b). The fact that PredNet develops very similar changes in encoding accuracy to those we measured in human observers suggests that these adaptive reallocations of coding resources indeed help to better predict future sensory input. It also demonstrates that predictive coding can implement a form of adaptation as proposed by our reallocation model.

**Figure 8:**
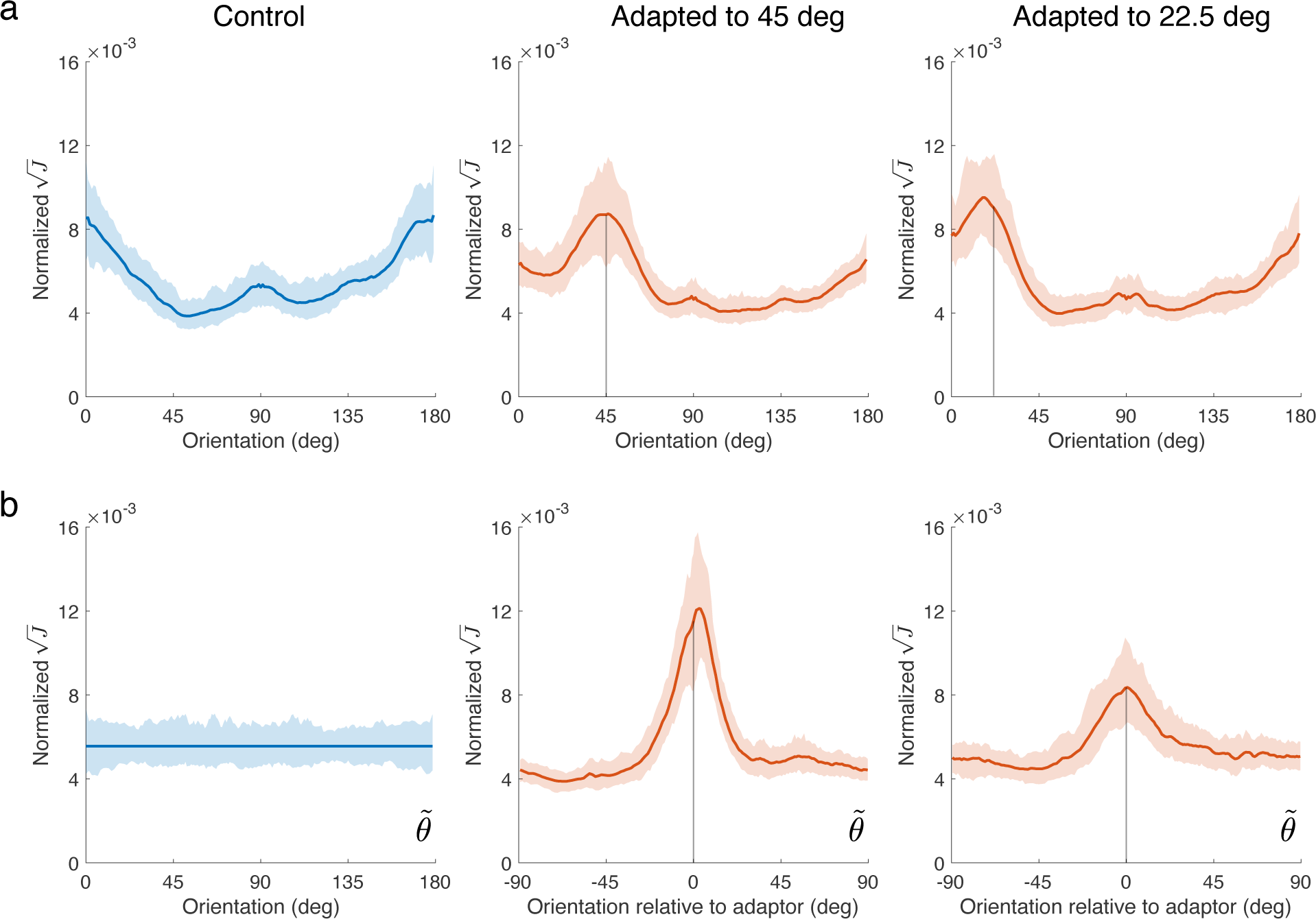
Encoding accuracy in the first representational layer *R*_0_ of PredNet after adaptation. (a) Normalized square root of Fisher information as a function of test orientation (0 deg is vertical). In the control adaptor condition (blue curve) Fisher information is higher at cardinal orientations, reflecting the fact that PredNet creates efficient representations of visual orientation given the predominance of cardinal orientations in natural scenes. After adaptation to a 45 deg (middle) or 22.5 deg (right) oblique adaptor, however, Fisher information peaks at the adaptor orientation. (b) Same as in (a) but plotted in sensory space 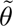. Similar to the encoding analysis of our psychophysical data, this sensory space is defined as the transformation 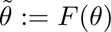 that leads to a uniform Fisher information distribution under the control adaptor condition (see also Fig. 1, Methods). When plotted relative to the adaptor orientation, Fisher information in sensory space is qualitatively similar to the adaptation kernels of human subjects (see Fig. 5a): Fisher information is peaked at and symmetric about the adaptor orientations, but reduced for test orientations further away. Lines and shaded areas represent the mean and the 95% confidence intervals over 200 stimulus sequences, respectively.

## Discussion

In this paper, we provide converging evidence that sensory adaptation in the human visual system adjusts encoding accuracy such that it is optimized for future sensory input. We psychophysically measured adaptation induced changes in sensory representation of visual orientation in human subjects. We found that these changes (relative to a non-oriented control adaptation state) are best described as a reallocation of sensory coding resources according to an isomorphic kernel that peaks at the adaptor orientation. Analyzing the temporal statistics of the retinal input of freely behaving human observers revealed that the distribution of local visual orientations in the next retinal input depends on the preceding stimulus history: the distribution shows a sharp peak at the mean orientation of a relatively stable stimulus history, but approaches a uniform distribution for histories with increasingly larger variances. This qualitatively matches the psychophysically measured changes in encoding accuracy, both for the control and the oblique adaptors. It suggests that adaption indeed ensures that sensory coding resources are efficiently allocated for future retinal input depending on the specific temporal structure of the input history. Finally, we asked whether, and if so how, a recurrent neural network that is optimized to predict the next frame of natural scene videos changes its sensory representations when being exposed to the same stimulus sequences used in our psychophysical adaptation experiments. We found that in the representational layers of the network, a reallocation of coding resources emerges that is very similar to that of human observers. This suggests that adaptation induced changes in sensory encoding are indeed aimed at allowing an optimal prediction of future sensory input.

Although previous studies have measured changes in discriminability as a result of adaption to a single oriented stimulus (Regan and Beverley, 1985; Clifford et al., 2001), none of them did so across the full range of orientations and against a well-defined control adaptor condition. These limitations have previously prohibited a comprehensive quantitative characterization of adaptation induced changes in sensory encoding. Interestingly, we found that adaptation not only decreases discrimination thresholds at but also orthogonal to the adaptor orientation, thereby confirming previous evidence (Clifford et al., 2001; Dragoi et al., 2002). A normative explanation of this orthogonal improvement remains unclear as neither the retinal input statistics nor the encoding changes in PredNet showed an effect at orthogonal orientations. We suspect that the effect is caused by the specific neural mechanisms involved in increasing encoding precision at the adaptor orientation (e.g., repulsive tuning curve shifts (Schwartz et al., 2007)). Furthermore, measuring the effects of adaptation relative to a well-defined control adaptor condition is crucial. Because adaptation affects the representation of every stimulus parameter (e.g. contrast, spatial frequency etc.), encoding changes superimpose throughout the processing hierarchy (Xu et al., 2008; Stocker and Simoncelli, 2009). The control adaptor was identical to the two oblique adaptors in every aspect except its orientation spectrum, allowing us to cleanly isolate the orientation specific, adaptation induced changes in orientation representation.

Temporal statistics of visual orientation in video sequences of natural environments have been previously investigated, although not at the level of the retina. For example, (Kayser et al., 2003; Betsch et al., 2004) analyzed the recordings taken by a camera mounted on a cat’s head. From the resulting video footage, they computed the correlation of orientation amplitudes over time at the same position for the same orientation, and showed that a prominent orientation is likely to repeat itself when the time interval is short while the correlation decreases as the time interval increases. Another study analyzed video footage taken with a static camera in a natural environment (van Bergen and Jehee, 2019). The distribution of the difference between the dominant orientations in successive frames showed that the next stimulus is more likely to be similar to than different from the previous one. Both of these results are qualitatively in agreement with our results and are not unexpected given the mostly continuous and temporally smooth statistics of natural environments (Schwartz et al., 2007; Gold and Stocker, 2017). However, human vision is based on active sensing (Yarbus, 1967; Hayhoe and Ballard, 2005; Findlay and Gilchrist, 2003; Najemnik and Geisler, 2005), where the active control of head and in particular eye-movements reshapes the statistical structure of the retinal input (Rothkopf et al., 2009; Rothkopf and Ballard, 2009). As a result, it has been unclear to what degree the spatiotemporal statistics of the natural environment translates to the statistics of the retinal image. Using a portable recording system that included a head-mounted camera in combination with mobile eye-tracking allowed us to characterize the statistical structure of the actual input to the human visual system under natural behavioral and environmental conditions.

A limitation of our current data set is that it mainly consists of forest scenes. While previous studies have found that the overall orientation statistics in natural scenes are slightly different depending on whether they contain man-made objects or not, as well as whether they are from indoor or outdoor environments (Coppola et al., 1998), it is unlikely that the conditioned temporal distributions we considering here are substantially different under different environmental contexts. Nonetheless, future studies are necessary to investigate this further (DuTell et al., 2022).

We demonstrate the functional benefit of sensory adaptation by analyzing the encoding dynamics of an artificial recurrent neural network (PredNet) trained to predict future sensory input. Artificial neural networks have become a powerful model to study normative explanations of neuronal and behavioral phenomena (Olshausen and Field, 1996; Singer et al., 2018). The rationale is that if a network, optimized for a certain functional goal within a certain stimulus environment, shows some emergent encoding characteristics, then they are likely beneficial for the network in achieving said functional goal in said environment. Previous studies have shown that sensory representations in artificial neural networks trained to identify objects in natural scenes are similarly shaped by the statistical context (i.e., priors) of the stimulus environment as the representations in the human visual system (Benjamin et al., 2019; Henderson and Serences, 2021). Here we now further demonstrate that this similarity extends to changes in these representations caused by the temporal context of typical adaptation experiments when the neural network is optimized fomaking predictions in a dynamic natural environment. This supports the efficient coding hypothesis. Note that PredNet does not contain adaptive neurons. Rather, the change in encoding emerges through the dynamic information flow in its recurrent architecture during the adaptation stimulus sequence. This shows that the functional goal of sensory adaptation is independent of a specific neural mechanism such as a reduction in response gain (Kohn and Movshon, 2003); neuronal gain changes may only be one of many ways to implement this goal.

The separation between function (i.e., encoding principle) and implementation (i.e., neural mechanisms) allows for a more general yet at the same time also more parsimonious definition of adaptation. Adaptation has been frequently characterized in terms of how it changes the tuning characteristics of individual and population of neurons. In addition to a reduction in neural gain, multiple effects such as changes and shifts in tuning curves (Dragoi et al., 2000; Kohn and Movshon, 2003) as well as homeostatic control of the neural populations activity have been attributed to adaptation (Benucci et al., 2013). However, these effects seem specific to certain cortical areas (Kohn and Movshon, 2004) and probably also certain animal species (Tring et al., 2023), and thus likely depend on the specific biophysical constraints and limitations. Here, by rather focusing on the changes in information capacity and how it relates to the future statistics of sensory input, we show that adaptation can be understood as following a single functional principle that, however, may rely on different specific neural implementations case by case. In that regard our results complement previous work on modeling adaptation as a trade-off between neural encoding costs and information loss (Młynarski and Hermundstad, 2021; Młynarski and Tkačik, 2022). Clearly, at the level of implementations these trade-offs may play an important role. Our results provide new insights into the functional objectives of such trade-off considerations.

It is worth discussing in more detail the idea of the (stationary) sensory space for which we characterize the adaptation kernel (Fig. 1); the kernel is isomorphic only in this space. The space reflects the efficient sensory representation of the stimulus feature given the overall, stationary stimulus statistics in the environment (Wei and Stocker, 2012, 2015). For visual orientation, for example, these statistics reflect the characteristic over-representation of cardinal orientations (Coppola et al., 1998). The representational geometry of the space (to use a currently popular term) is such that the statistics and the encoding precision are homogeneous and uniform. We propose to think of adaptation as a transient modulation or fine-tuning of this geometry such that it is optimally prepared for the specific stimulus context induced by the short-term stimulus history.

This view also allows us to think of adaptation with regard to more generally construed “statistical” contexts. For example, it is well documented that spatial context can change sensory encoding of visual orientation in similar ways as temporal context does, both at the neural and behavioral level (Schwartz et al., 2007, 2009). The changes are also well-aligned with the fact that in natural scenes, visual orientation at a location is best predicted by the average orientation within its surround (Felsen et al., 2005). Furthermore, these spatial contexts are specific to the particular image content, and the associated modulation of neural response patterns can be explained with a flexible gating mechanisms that is optimally tuned for these specific contexts (Coen-Cagli et al., 2015). Thus we conjecture that sensory adaptation is but one mechanism by which neural systems ensure sure that their representations of sensory information is efficient with regard to the statistical structure of their environment.

## Conclusions

Efficient coding has been a prominent hypothesis for sensory adaptation. The present study mapped out the change in coding accuracy across the feature space after adapting to a single feature value through a psychophysical experiment, and found a universal parametric description of the reallocation of coding resources. Analysis of the temporal statistical structure of retinal images in freely behaving humans and sensory representations in recurrent neural networks provide converging support for the efficient coding hypothesis of adaptation: adaptation induced changes in encoding accuracy and perceptual behavior that reflect the visual systems’ attempt to best possibly represent the next expected sensory input.

## Acknowledgments

The authors thank the members of the Computational Perception and Cognition Laboratory for many fruitful discussions and feedback. The presented research was supported by the University of Pennsylvania. JM was in part supported by the National Science Foundation and DoD OUSD (R&E) under cooperative agreement PHY-2229929 (The NSF AI Institute for Artificial and Natural Intelligence). Some of the results described in this paper have been presented at the Annual Meeting of the Vision Science Society in May 2023.

## Methods

### Psychophysical experiment

#### Subjects

5 subjects (2 female), 25 to 33 years old, participated in the experiment. Subject 1 was non-naive. All subjects had normal or corrected-to-normal vision. All subjects gave written consent to participate under the condition that they could quit at any moment. They were remunerated at a rate of 10$ per hour, plus a 20$ bonus upon completion of the full experiment.

#### Setup

The experiment was run using Matlab (R2016b) with the PsychToolbox (Brainard, 1997). Subjects sat in a darkened room and viewed stimuli on a VPixx3D screen (1920×1080 pixels resolution, 120 Hz refresh rate) at a 89 cm distance. A circular aperture (26 cm diameter) was placed in front of the screen to occlude the edges of the screen removing any potential cardinal orientation cues.

#### Stimuli

All stimuli were presented on a gray background with mean luminance 40 cd/m^2^. Stimuli consisted of filtered white noise patterns with same overall mean luminance as the background. For all stimuli (control and oblique adaptors, test), the noise was first filtered with a band-pass filter with uniform power spectrum across the spatial frequency range of 3.75 – 5.25 cpd. Oblique adaptors and test stimuli were then further filtered with an oriented filter with a symmetrically warped Laplace profile centered at the desired orientation and a standard deviation of 1.4 deg. Stimuli had 80% contrast. Stimuli were 2 deg in diameter, presented 1.67 deg to the left and right of fixation. A fixation dot was presented at the center of the screen throughout the experiment.

#### Procedure

The experiment was organized in blocks (see below). At the beginning of a block, subjects viewed one of the adaptor stimuli for 60s. After this initial adaptation phase, each trial started with top-up adaptation (5s), followed by a blank interval (0.35s), presentation of the test screen (0.1s), and a response period. During the initial adaptation phase and the top-up period, two identical adaptor patterns were presented on both sides of fixation, refreshing every 1.25s with a 0.05s blank frame in between. In the test screen, a test and a reference stimulus were presented to the left and right of fixation, randomly assigned. During the response period, subjects pressed one of two buttons on a gamepad to indicate which stimulus in the test screen was more clockwise (or counterclockwise, interleaved across blocks). Subjects were instructed to maintain fixation throughout the initial adaptation phase and during each trial.

The experiment consisted of two parts, each of which contained an oblique adaptor condition and a control adaptor condition. In the first half of the experiment, the oblique adaptor was oriented at *±*45 deg. Vertical corresponds to 0 deg. The test orientations were [0, *±*10, *±*30, *±*45, 90] deg relative to the adaptor orientation (subject 1 had *±*5 deg in addition). In the second half of the experiment, the oblique adaptor was oriented at *±*22.5 deg. The test orientations were [0, *±*10, *±*22.5, *±*45, 90] deg relative to the adaptor orientation. Test orientations were randomized across trials. The reference orientation varied according to a 2-up-1-down staircase procedure in 25 equal steps within a *±*9.6 deg, *±*15 deg, *±*18 deg, or *±*24 deg range relative to the test orientation, depending on the performance of each subject in the training session prior to the experiment.

Subjects completed 192 trials for each test orientation in each adaptation condition (216 trials for subject 1 in the first half of the experiment). In each half of the experiment, subjects completed the control adaptor condition first, oblique adaptor condition next. In the oblique adaptor condition, subjects completed 4 blocks of one adaptor orientation, then 4 blocks of the same adaptor orientation but mirrored around vertical (6 blocks each for subject 1 in the first half), e.g. first 45 deg then −45 deg. The order of the two adaptor orientations were counterbalanced across subjects. The control adaptor condition consisted of 8 blocks (12 blocks for subject 1 in the first half). Each block lasted for about 25 min, depending on the response time of each subject. Blocks with different adaptors (control versus oblique adaptor condition, or opposite oblique adaptor orientations) were completed at least one day apart.

### Data analysis

For the main analysis, data in blocks with −45 or −22.5 deg oblique adaptors were combined with data from blocks with 45 or 22.5 deg adaptor by mirroring it across vertical. Psychometric curves were obtained by fitting cumulative Gaussian distributions to the data. We assumed a mean of 0 deg for the Gaussian distribution and zero lapse rate. Discrimination thresholds were calculated at the 75% level based on the fitted psychometric functions.

### Modeling

#### Pre-adaptation (control adaptor)

Let *θ* be the orientation of the test stimulus and *m* its sensory measurement in a given trial. Before adaptation (i.e., defined by the control adaptor condition) the discrimination threshold is typically lower at cardinal orientations (Caelli et al., 1983), which implies higher Fisher information at cardinal orientations (Wei and Stocker, 2015). We parametrized the square root of the Fisher information distribution *J* (*θ*) as the weighted sum of a uniform distribution and two identical von Mises distributions centered at two cardinal orientations:

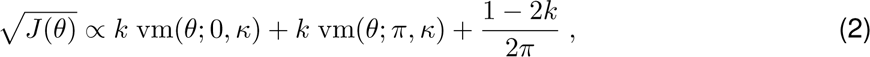

where *κ* represents the width of the distribution around cardinal orientations, and *k* represents the relative amplitude. Note that because angles are defined on [*−π, π*], 0 and *±π* represents cardinal orientations (0 corresponds to vertical). We use this convention throughout the paper.

We consider encoding under the control condition as reflected by a sensory space (stationary) in which Fisher information is uniform. The mapping 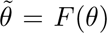 from stimulus to this sensory space is the cumulative of the square root of Fisher information distribution, thus 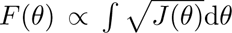. Assuming that sensory noise in this space follows a von Mises distribution, we can write the measurement distribution in sensory space as

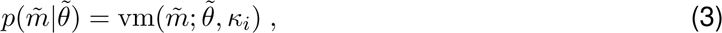

where *κ_i_*represents the sensory noise magnitude. The distribution in stimulus space *p*(*m|θ*) directly follows by transformation according to the inverse mapping 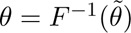.

#### Post-adaptation (oblique adaptors)

After adapting to a single orientation *θ_a_*, the distribution of the square root of Fisher information in sensory space changes accordingly to

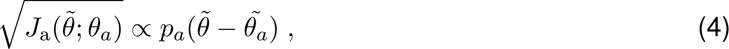

where 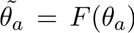 is the adaptor orientation in sensory space, and *p_a_* is the adaptation kernel. Thus, adaptation reallocates Fisher information by the same adaptation kernel shifted according to the adaptor. Now, the stationary sensory space is <u>not a</u> uniform space any more. However, we can apply another transformation 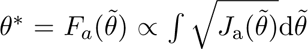 to obtain an adapted sensory space.

Importantly, we assume that in the transformed space sensory noise remains von Mises distributed with the same internal noise parameter as in the stationary sensory space (Eq. 3), thus the total coding resource does not change after adaptation (reallocation). We can write the measurement distribution as

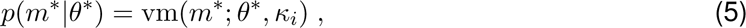

where *κ_i_* represents the constant sensory noise magnitude. The distribution in stimulus space *p*(*m|θ*) then follows by successive transformations according to the inverse mappings 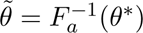 and 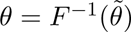.

The “2-peak” (original) model assumes that the adaptation kernel 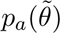 is a weighted sum of two independent von Mises distributions and a uniform distribution:

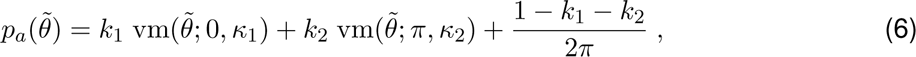

where *k*_1_ and *k*_2_ represent the relative amplitudes, and *κ*_1_ and *κ*_2_ the widths of the two peaks.

The “1-peak” model assumes that 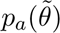 is the weighted sum of one von Mises distribution and a uniform distribution:

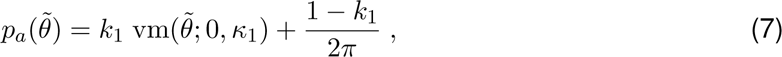

where *k*_1_ and *κ*_1_ represents the relative amplitude and the width of the peak, respectively.

The “2-peak + Fisher” model assumes that total Fisher information can change after adaptation, i.e., the sensory noise (i.e. the width of the von Mises likelihood) in the adapted sensory space 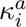 is allowed to be different from the the noise in the stationary-adaptation sensory space *κ_i_*.

The “2-peak + kernel” model permits that adaptation kernels 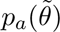 for the 45 deg and 22.5 deg adaptor condition can be different.

#### Discrimination decision and response distribution

Let *θ_t_* and *θ_r_* be the orientation of the test and reference stimulus, and *m_t_* and *m_r_*their sensory measurements respectively. The probability of the reference orientation being more clockwise than the test orientation can be calculated as

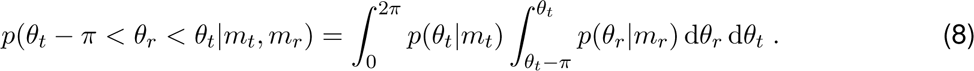

If the probability is larger than 0.5, the observer would make the decision that the reference orientation is more clockwise; otherwise the test orientation.

Because test and reference stimuli were always tested for the same adaptation state of the observer, had the same probability to be clock-wise or counter-clockwise, and the stimuli had the same spatial frequency, contrast and presentation duration, the decision process can be simplified to a direct comparison of the measurements of the two stimuli: if *m_r_* is more clockwise than *m_t_*, the subjects would make the decision that the reference orientation is more clockwise, and vice versa. So the decision probability of the reference stimulus being more clockwise can be written in terms of the measurement distributions (Eq. (3) and (5) transformed to stimulus space), thus

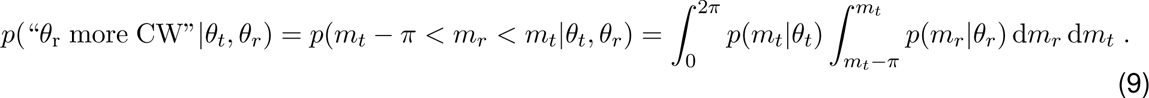

### Model fitting

We fit the model by maximizing the likelihood of the data given the model:

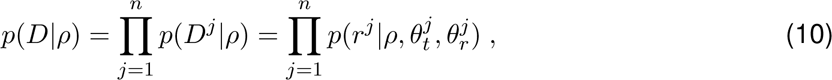

where *D* is the data, *ρ* represents the parameters of the model, 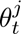 and 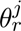 are the test and reference orientations and *r^j^*is the response in trial *j*, and *n* is the total number of trials.

We first fit the model to the control adaptor condition with the following free parameters:

- *κ_i_* for sensory noise;
- *k* for the relative peak amplitude, and
- *κ* for the width of the Fisher information distribution (von Mises).

Then we fix these parameters and fit the model to the oblique adaptor condition.

The 2-peak model has four free parameters for the adaptation kernel:

- *k*_1_ and *k*_2_ for the relative amplitudes of the two peaks;
- *κ*_1_ and *κ*_2_ for the width of the two von Mises distribution.

The 2-peak + Fisher model has an additional parameter 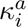 allowing a different sensory noise after adapting to an oblique adaptor (i.e. change in overall coding resource). The 2-peak + kernel model has four parameters for the adaptation kernel of each oblique adaptor, eight in total. The 1-peak model has only two free parameters for the strength and width of the adaptation kernel.

### Natural scene statistics

#### Data set

The dataset was obtained with a video-based, portable eye tracker system (Eye Tracking Glasses 2 from SensoMotoric Instruments). Videos were filmed by the head-mounted camera while participants were freely walking in a wooded environment. Image sequences had a resolution of 24 fps, 1280×960 pixels, spanning 60×46 deg of visual angle. The compression algorithm was H.264. Synchronized eye movements were recorded binocularly at 120 samples per second. Initial calibration of each participant was accomplished with a 3 point calibration target and participants subsequently wore the eye tracker for at least 10 minutes before checking the calibration again. The resulting eye tracking accuracy was well below 1 degree at about 10m distance. We included videos from 9 participants, with a total length of 12 minutes. Videos were converted to grayscale for analysis.

#### Data analysis

We looked at a 6×6 deg area centered at the fixation in each frame of the videos. We extracted the orientation at each position within the area using a steerable pyramid decomposition (Simoncelli and Freeman, 1995). We rotated and applied the 1st-order steerable filter to find the orientation with the strongest response as the orientation of each position. We then computed the mean and circular variance of orientation over 3s time-windows (72 frames) at each position, and computed the difference between the orientation in the next frame and the mean orientation in the previous 3s. We included the results from level 2 to 4 of the steerable pyramid. In Fig. 6, we included time and positions where the history variance is smaller than 0.1 or larger than 0.9, respectively, and combined the data from three levels.

### PredNet

PredNet is a recurrent neural network that predicts the next frame of a video. We used PredNet pretrained on the KITTI data set for our experiment (Geiger et al., 2013; Lotter et al., 2016).

#### Stimuli

The stimuli were images of filtered white noise patterns with a size of 128×160 pixels. The control adaptors were filtered by the spatial frequency spectrum extracted from the training data set within the range of 8-12 cycles per image. The spatial frequency filter was obtained by taking the average of the 2D Fourier transformation of the image across all frames and averaging across orientation for each spatial frequency, with a low and high cutoff at 8 and 12 cycles per image respectively. The oblique adaptors and test stimuli were further filtered by an orientation filter with a symmetrically wrapped Laplace spectrum centered at the desired orientation with a standard deviation of 1.4 deg, in addition to the same spatial frequency filter as the control adaptor. Stimuli had 100% contrast. The noise pattern was embedded in a circular aperture at the center of the image; the contrast of the noise pattern fades linearly from 100% to 0 as the distance from the center increases from 48 to 60 pixels.

#### Procedure

Each input sequence consisted of four adaptor frames followed by a test frame. We fed this five-frame input sequence to PredNet and extracted the activation of the first representational layer in response to the test frame (frame #5). For each adaptation condition (control, oblique 22.5 and 45 deg), we tested 200 input sequences (different noise patterns, exact same filter properties). To compute Fisher information as a function of test orientation, we rotated the test frame in each sequence in 1 deg intervals.

#### Calculating Fisher information

We computed Fisher information in the first representation layer (*R*_0_) as a function of orientation *θ* in the test frame (Fig. 7a). Assuming independent Gaussian noise, Fisher information can be calculated for each test frame as

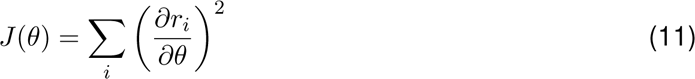

where *r_i_* is the response of the *i*th unit in layer *R*_0_. Because we focus on the distribution of coding resources, we normalized the sum of the square root of Fisher information across orientation. Figure 8 shows the mean and 95% confidence intervals over the 200 input sequences.

## Supplementary information

**Supplementary Table 1:** Best-fitting model parameters for every subject.

| Parameter | Subj 1 | Subj 2 | Subj 3 | Subj 4 | Subj 5 |
| --- | --- | --- | --- | --- | --- |
| **Control** |  |  |  |  |  |
| $\kappa_i$: sensory noise | 191.04 | 128.63 | 25.44 | 60.54 | 128.46 |
| $k$: prior intensity | 0.17 | 0.24 | 0.12 | 0.25 | 0.15 |
| $\kappa$: prior width | 14.73 | 22.47 | 15.49 | 14.69 | 16.83 |
| **2-peak** |  |  |  |  |  |
| $k_1$: kernel strength | 0.05 | 0.08 | 0.09 | 0.11 | 0.08 |
| $\kappa_1$: kernel width | 74.8 | 77.2 | 39.7 | 31.8 | 105.7 |
| $k_2$: kernel strength | 0.13 | 0.11 | 49.47 | 0.25 | 49.49 |
| $\kappa_2$: kernel width | 3.17 | 2.32 | 0.0046 | 2.17 | 0.0071 |

**Supplementary Table 2:** Best-fitting model parameters for every subject - alternative models.

| Parameter | Subj 1 | Subj 2 | Subj 3 | Subj 4 | Subj 5 |
| --- | --- | --- | --- | --- | --- |
| **1-peak** |  |  |  |  |  |
| $k_1$: kernel strength | 0.04 | 0.07 | 0.07 | 0.07 | 0.06 |
| $\kappa_1$: kernel width | 127.2 | 93.4 | 64.2 | 77.0 | 182.8 |
| **2-peak + Fisher** |  |  |  |  |  |
| $\kappa_i^a$: sensory noise (adapt) | 185.08 | 143.58 | 25.43 | 48.33 | 108.98 |
| $k_1$: kernel strength | 0.05 | 0.08 | 0.09 | 0.12 | 0.08 |
| $\kappa_1$: kernel width | 74.4 | 78.4 | 39.7 | 30.4 | 108.9 |
| $k_2$: kernel strength | 0.12 | 0.17 | 47.23 | 0.18 | 1.15 |
| $\kappa_2$: kernel width | 3.53 | 1.52 | 0.0048 | 2.45 | 0.27 |
| **2-peak + kernel** |  |  |  |  |  |
| $k_1^{45}$: 45° kernel strength | 0.05 | 0.09 | 0.07 | 0.09 | 0.08 |
| $\kappa_1^{45}$: 45° kernel width | 134.6 | 75.2 | 92.0 | 49.2 | 92.3 |
| $k_2^{45}$: 45° kernel strength | 0.13 | 0.01 | 0.09 | 0.21 | 0.39 |
| $\kappa_2^{45}$: 45° kernel width | 5.21 | 700.0 | 7.53 | 7.52 | 1.22 |
| $k_1^{22.5}$: 22.5° kernel strength | 0.06 | 0.07 | 0.12 | 0.09 | 0.07 |
| $\kappa_1^{22.5}$: 22.5° kernel width | 35.8 | 79.1 | 22.9 | 46.4 | 191.9 |
| $k_2^{22.5}$: 22.5° kernel strength | 0.12 | 0.19 | 415 | 0.35 | 381 |
| $\kappa_2^{22.5}$: 22.5° kernel width | 1.96 | 1.27 | 5.7e-4 | 0.71 | 7.6e-4 |

**Supplementary Figure 1:**
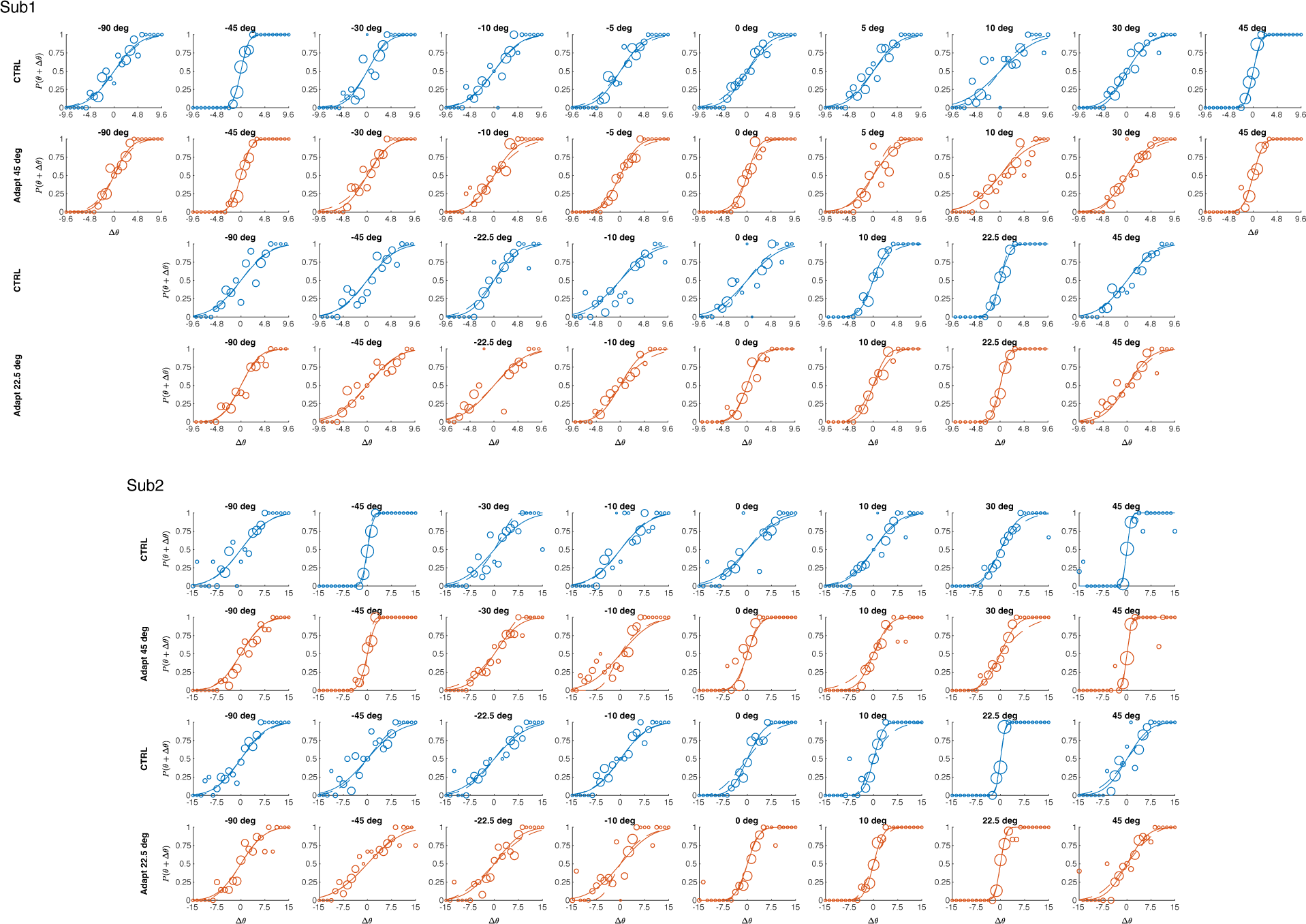
2AFC response data of subjects 1 & 2 and psychometric curves fitted by a cumulative Gaussian distribution (solid line) or the reallocation 2-peak model (dashed line). Size of the data point represents the number of trials. Angle listed above each subplot indicates the difference between the test and the adaptor orientation.

**Supplementary Figure 2:**
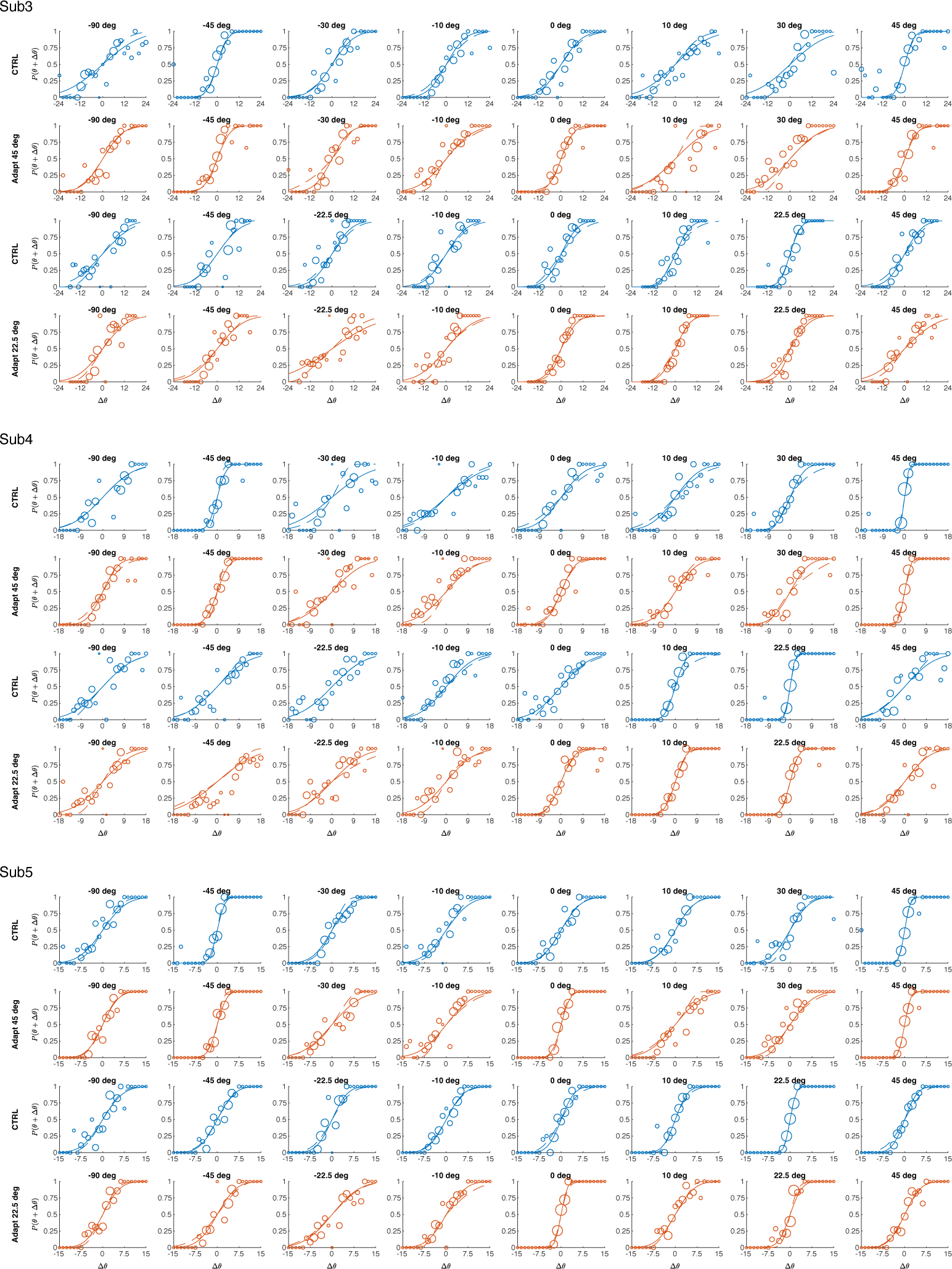
2AFC response data of subjects 3, 4 & 5 and psychometric curves fitted by a cumulative Gaussian distribution (solid line) or the reallocation 2-peak model (dashed line). Size of the data point represents the number of trials. Angle listed above each subplot indicates the difference between the test and the adaptor orientation.

**Supplementary Figure 3:**
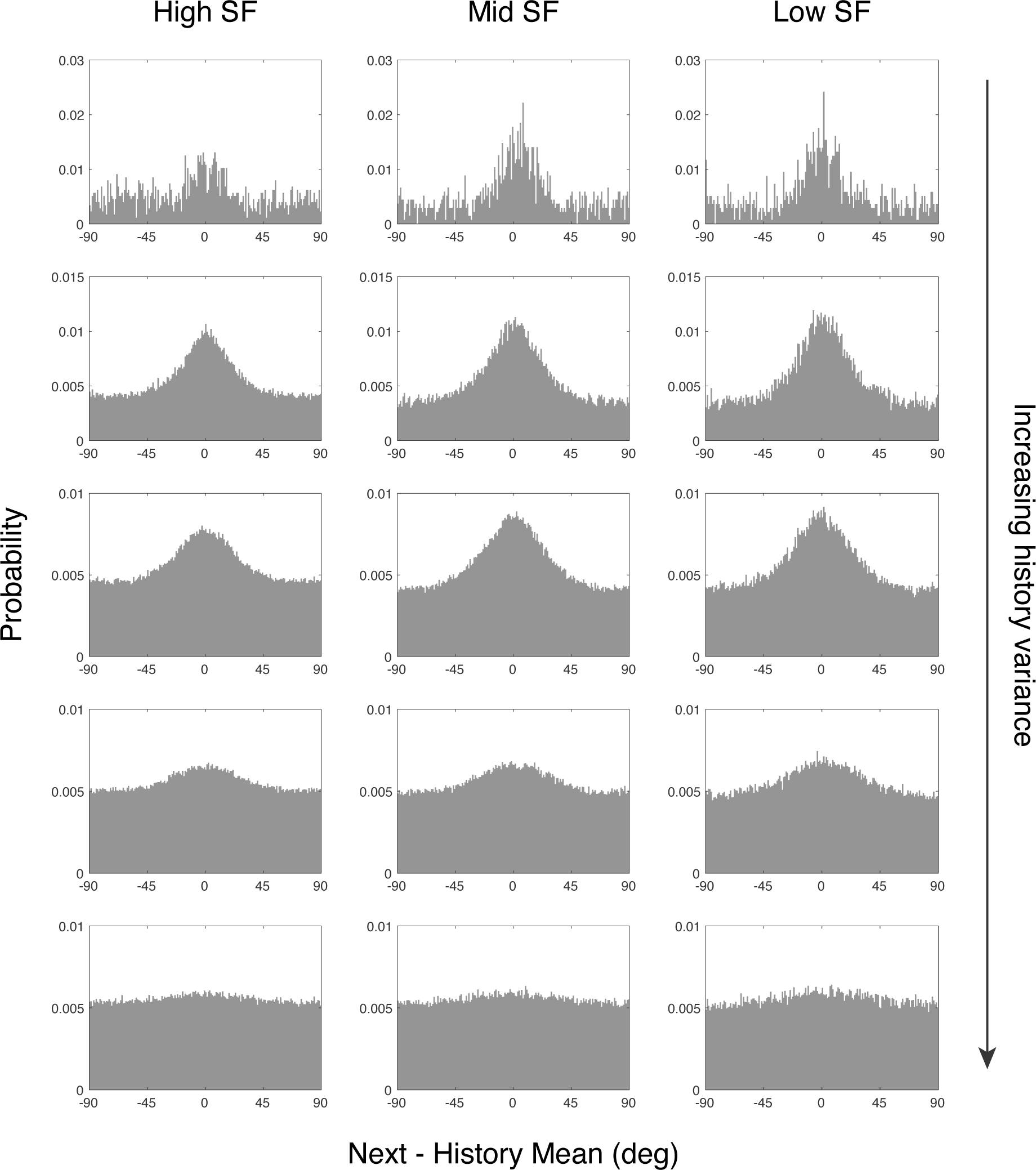
Distribution of orientation in the next frame relative to history mean (across 3s time-window) for different spatial frequencies and different history variance. The three spatial frequency levels were calculated from level 2 to 4 of the steerable pyramid and approximately correspond to 2.5, 1.25, and 0.63 cycles/deg, respectively. Variance are circular variance ranging from 0 to 1 and binned into bins of 0.2. For all spatial frequency levels, the distribution of orientation in the next frame is more concentrated around the history mean as the variance becomes smaller.

